# Counting is Almost All You Need

**DOI:** 10.1101/2022.08.09.501430

**Authors:** Ofek Akerman, Haim Isakov, Reut Levi, Vladimir Psevkin, Yoram Louzoun

## Abstract

The immune memory repertoire encodes the history of present and past infections and immunological attributes of the individual. As such, multiple methods were proposed to use T-cell receptor (TCR) repertoires to detect disease history. We here show that the counting method outperforms all existing algorithms. We then show that the counting can be further improved using a novel attention model to weight the different TCRs. The attention model is based on the projection of TCRs using a Variational AutoEncoder (VAE). Both counting and attention algorithms predict better than any current algorithm whether the host had CMV and its HLA alleles. As an intermediate solution between the complex attention model and the very simple counting model, we propose a new Graph Convolutional Network approach that obtains the accuracy of the attention model and the simplicity of the counting model. The code for the models used in the paper are provided in: https://github.com/louzounlab/CountingIsAlmostAllYouNeed

## Introduction

Following recent developments in immune sequencing technology (18; 7; 3), large T-Cell Receptor (TCR) repertoires can be sampled. Given the association of diseases and TCRs, such repertoires could in theory be used for systemic detection of disease history. However, methods to decipher the disease history from these repertoires (currently denoted “reading the repertoire”) are still limited. Recently, Bayesian approaches and machine learning methods to read repertoires (15; 33; 42; 39; 52; 55) were proposed in this field, with a good accuracy. However, even those do not reach the accuracy required for clinical usage.

From a computational point of view, the repertoire classification problem is a Multiple Instance Learning (MIL) task. MIL problems arise when the training examples are of varying sizes. In MIL problems, a set or bag is labeled instead of a single object. In the standard definition, a bag *X* = {*x_i_*} receives a label *Y_X_* = *max*{*y_i_*} where *y_i_* is the label of *x_i_*. Here, *y_i_* ∈ {0, 1}. However, this can be extended to any label. During training, we are unaware of *y_i_*. Only *Y_X_*, the class of each bag in the training set, is known. Examples of MIL problems are video classification, where each frame is an instance, text classification, where each word is an instance, 3D object classification, where each point is an instance, and more (8; 48).

The standard MIL assumption can be expanded to address tasks where positive bags cannot be identified by a single instance. However, the bag can still be classified by the distribution, interaction or accumulation of the instances in the bag (8).

To formulate the TCR repertoire classification task as an MIL task, a repertoire can be viewed as a bag of TCR sequences, of which a very small fraction is associated with the class of interest. We use the following notations in the current analysis: *T* = {*t*_1_, *t*_2_, *t*_3_,…, *t_R_*} is the group of all TCRs in all samples (training or test) that may be very large. *X_j_* = {*t*_*j*_1__, *t*_*j*_2__, *t*_*j*_3__,…, *t*_*j_N_*_} is a specific repertoire and *Y*(*X_j_*) ∈ {0, 1} is the binary label of the repertoire *X_j_*. We further assume for the sake of notation simplicity that a TCR *t* can either bind or not bind any peptide *p*, with some arbitrary binding cutoff. We denote the set of TCR that binds the peptide *p* by *T*(*p*).

The TCR repertoire classification problem includes unique difficulties compared with classical MIL problems:

- **Low overlap** - The immune repertoire overlap of different individuals is low ((24; 14)). Given two repertoires *X_j_*, *X_k_*, |*X_j_* ∩ *X_k_*| is very small.
- **Non-injectivity of TCR-peptide binding** - Multiple sequences can bind to the same pathogen (53). |*T*(*p*)| > 1 for most target peptides.
- **Large TCR diversity** - Recent studies suggest that the human body can have *>* 10^14^ unique TCR sequences (36). |*T*| ≥ 10^14^.
- **An extremely low Witness Rate (WR)** - In MIL problems, the WR is defined by the percentage of discriminating instances within a bag. A WR of 1-5% is considered low in MIL tasks (52). We analyze here a large CMV binding dataset, used by multiple groups (52; 41; 14; 12). Each immune repertoire in the dataset has an average of 192,515 (±80,630 s.d) unique TCR sequences (15), of which we further estimate only an order of 100 are associated with CMV (15; 9), i.e., the WR can be lower than 0.0001%. Formally, for each repertoire *X_j_* and target peptide *p* 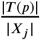 is very small.

We here show that counting arguments actually produce better results than the current SOTA ML or Bayesian methods. We then further improve on that by including the similarity between different TCRs using the combination of a Variational AutoEncoder (VAE) (13), and a novel attention model to include not only the relative importance of positive samples, but also their quantity, named attTCR (attention TCR). Finally, we propose an intermediate solution between the counting and attTCR - gTCR that uses a graph of the TCR repertoire co-occurences to predict the class of a sample.

### Related work

In recent years, ML and statistical data analysis tools have been proposed to solve the repertoire classification problem. Emerson et al (15) released a dataset composed of 786 immune repertoires, most of them with a CMV negative/positive classification as well as low resolution class-I HLA typing (for a detailed data description see the ‘Data’ section). They use a Fisher exact test to score TCRs based on their association with positive and negative repertoires, and classify TCR repertoires as either positive or negative to CMV or for a given HLA allele.

Their work has been enlarged by TCR-L (33) who evaluate the association between the TCR repertoire and clinical phenotypes. TCR-L expands on Emerson and also uses information about the structure of the TCR sequences and other information about the patient.

Machine learning models, and specifically attention based machine learning models, were also proposed as immune repertoire classifiers. deepTCR(42) implements multiple deep learning methods, and a basic form of attention-based averaging. deepTCR encodes each TCR*β* chain with a combination of its *V_β_*, *D_β_* and *J_β_* genes using a Convolutional Neural Network (CNN) that extracts sequence motifs. This information is further encoded using a VAE. Then, an attention score is given to each TCR using a custom attention function they designed called AISRU. Finally, a fully connected network (FCN) classifier determines the immune repertoire’s status.

Another recently developed model (39) uses 4-mers, sub-sequences of the TCRs CDR3. A logistic regression model is trained on the 4-mers as inputs. Similarly, MotifBoost (28) uses 3-mers to classify the repertoire, using GBDT (gradient boosted decision trees).

Finally, Deep-RC (52) implements an attention model and uses 1D CNNs in orderto embed every TCR to a fixed dimension. Those embeddings are forwarded to more FCN layers, and awarded attention scores using a Transformer-like (49) attention equation.

### Novelty

The algorithms presented here present multiple novel aspects to improve the accuracy of repertoire association studies.

First, we show that a simple counting argument obtains a higher accuracy than all previous methods.

We then propose a novel attention methods that on the one hand gives a different importance to different components, but on the other hand counts them. This is obtained through the sum over the attention of each TCR, with no softmax, but with sigmoid. We show that in contrast with classical attention models, the attention scoring with non-constant sum improves performance over the simple counting algorithm. The only normalization performed is on the sum of the attention scores, to put it in the active range for the loss function.

Finally, we combine the counting and attention in the Graph Neural Network (GNN) based gTCR model. We use a GNN to classify the repertoire. To the best of our knowledge, this is the first usage of GNN in TCR repertoire classification. The proposed GNN has two novel methodological aspects. First, the contribution of self edges in the modified adjacency matrix is learnt with the weights. Second, we use vertex identity aware graph classification. The combination of these two methods obtain the accuracy of the attention model with the simplicity of the counting one.

At the technical level, attTCR offers several improvements over Deep-RC (52) and deepTCR (42). The embedding method of each TCR using a cyclic variational autoencoder has never been used on TCRs.

The combination of these methods produce three levels of complexity for the model, where even the simplest model is more accurate than current state of the art (SOTA) models.

## Results

### Positive selection and detection of TCRs associated with a condition

Although, the TCR repertoire is very diverse, with most positions along the CDR3 highly variable (36; 4), still a large number of TCRs are shared among multiple patients.

We computed sharing of TCRs between samples in the Emerson dataset (15) (further denoted ECD), where a TCR is defined as the combination of *Vβ*, and *Jβ* genes and a CDR3 amino acid sequence (even with different nucleotide sequence). While most of the TCR sequences appear in a single repertoires, there are ~ 10^5^ unique TCRs that appear in more than 10 different repertoires, and hundreds of TCRs that appear in more than a 100 repertoires (Figure 1A). As such, there is enough intersection between different TCRs to perform classification algorithms.

**Figure 1.**
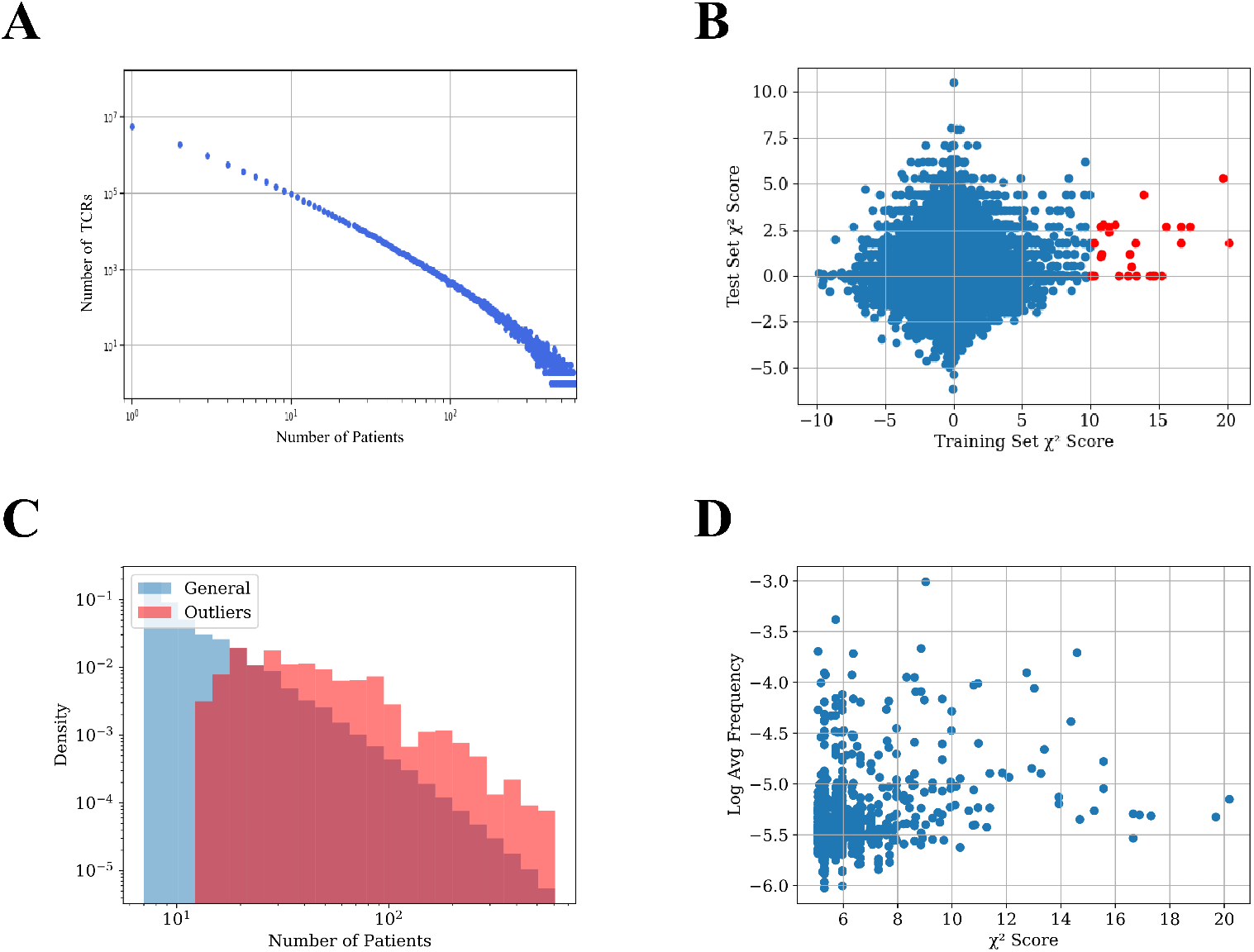
(A) TCR number as a function of the number of the patient repertoires that have them in the training set. (B) Distribution of the TCRs’ *χ*^2^ scores in the training and test sets. The x-axis value is the *χ*^2^ score of the TCR on the training set, the y-axis value is the *χ*^2^ score of the same TCRs on the test set. TCRs with an absolute *χ*^2^ score of over 10 in the training set are colored red. Notice that there are only such points on the positive side of the axis.(C) Distribution of average frequency per sample reactive and general TCRs in the dataset. General TCRs refer to all the TCRs in the dataset included in at least 7 repertoires, and reactive TCRs refer to the 200 TCRs with the highest *χ*^2^ score. The distribution of reactive receptors is clearly shifted to the right. (D) Scatter plot of different TCRs in the dataset. The x-axis represents the *χ*^2^ score of each TCR, and the y-axis represents its log average frequency in the repertoires it appears in. No correlation is observed between the two.

One can assume that following T cell clonal expansion, TCRs that bind to specific diseases are more frequent, and as such are likely to appear in repertoires of people who are or were infected by the disease. However, while we expect some TCRs to be positively associated with a disease or a condition, there is no a-priori reason for any TCR to be negatively associated with a condition (i.e., that its absence is evidence for a condition). To test the absence of negative selection by pathogen, we split the data into a training and a test set (see’Experimental setup’), and calculated the *χ*^2^ score between the expected and observed number of CMV positive patient that carry a TCR for both the train and test sets (see section ‘*χ*^2^’). We then multiplied the score by the sign of the difference of the expected and observed number of CMV+ patients carrying the TCR (i.e., TCRs less present in positive samples than expected have a negative sign - Figure 1B).

For the vast majority of the TCRs, the *χ*^2^ score is distributed around 0. However, there are some outliers with high *χ*^2^ scores in the training set. Many of those also have a high *χ*^2^ score in the test set (red points). More interestingly, the deviation is only on the positive side. In other words, some TCRs are strongly positively associated with the CMV+ patient class. However, as expected, there are no TCRs associated with the CMV-patient class. We propose to use (only) the TCRs positively associated with the condition (CMV in this case) in the training set to classify patients.

### No systemic difference between CMV+ and CMV-samples

High *χ*^2^ score reactive TCRs are obviously more likely to be shared between more repertoires than the other TCRs (Figure 1C), since a non-shared receptor per definition has a low *χ*^2^ score. Although reactive TCRs go through clonal expansion, checking which TCRs have a large frequency within the repertoire of each donor is not a sufficient method to find such reactive TCRs. Figure 1D demonstrates the lack of correlation between the *χ*^2^ score of each TCR and its average frequency in the samples where it is present.

Instead of focusing on a specific TCR, one could propose to use more generic features of the repertoire to distinguish between CMV+ and CMV-patients ((46; 23; 43)). This may be true for lytic conditions, but not for latent or historical conditions. We expect no difference in the general properties of the peripheral repertoire. For events in the distant past, most of the TCRs that were active during the immune response are no longer in the blood in high quantities, and when looking at the general data distribution in the repertoire, there is no difference between positive and negative repertoires (see the Appendix for comparison between V, and J gene distributions and the CDR3 compositions of CMV+ and CMV- patients).

### Counting is all you need

Given the association of specific TCRs with a condition, one could propose different methods to combine reactive TCRs into a classifier for disease history. We here argue that counting the number of such TCRs in a repertoire is a better classifier than any existing complex ML classifier.

To clarify that, we propose a simplistic model that captures the essence of the problem. Assume a general very large set of TCRs, where each patient has a random subset of these TCRs. Within the large set of TCRs, there is a small subset associated with the disease, and patients that had the disease have a higher than random chance of having these TCRs (see Figure 2 for a description of the model). The data generation process uses 3 probabilities: *p*_0_ - the probability that a TCR would be selected in any patient, *p*_1_, *p*_2_ - the probability that a selected TCR associated with CMV is added to a repetoire in CMV positive and negative samples (Fig. 2).

**Figure 2.**
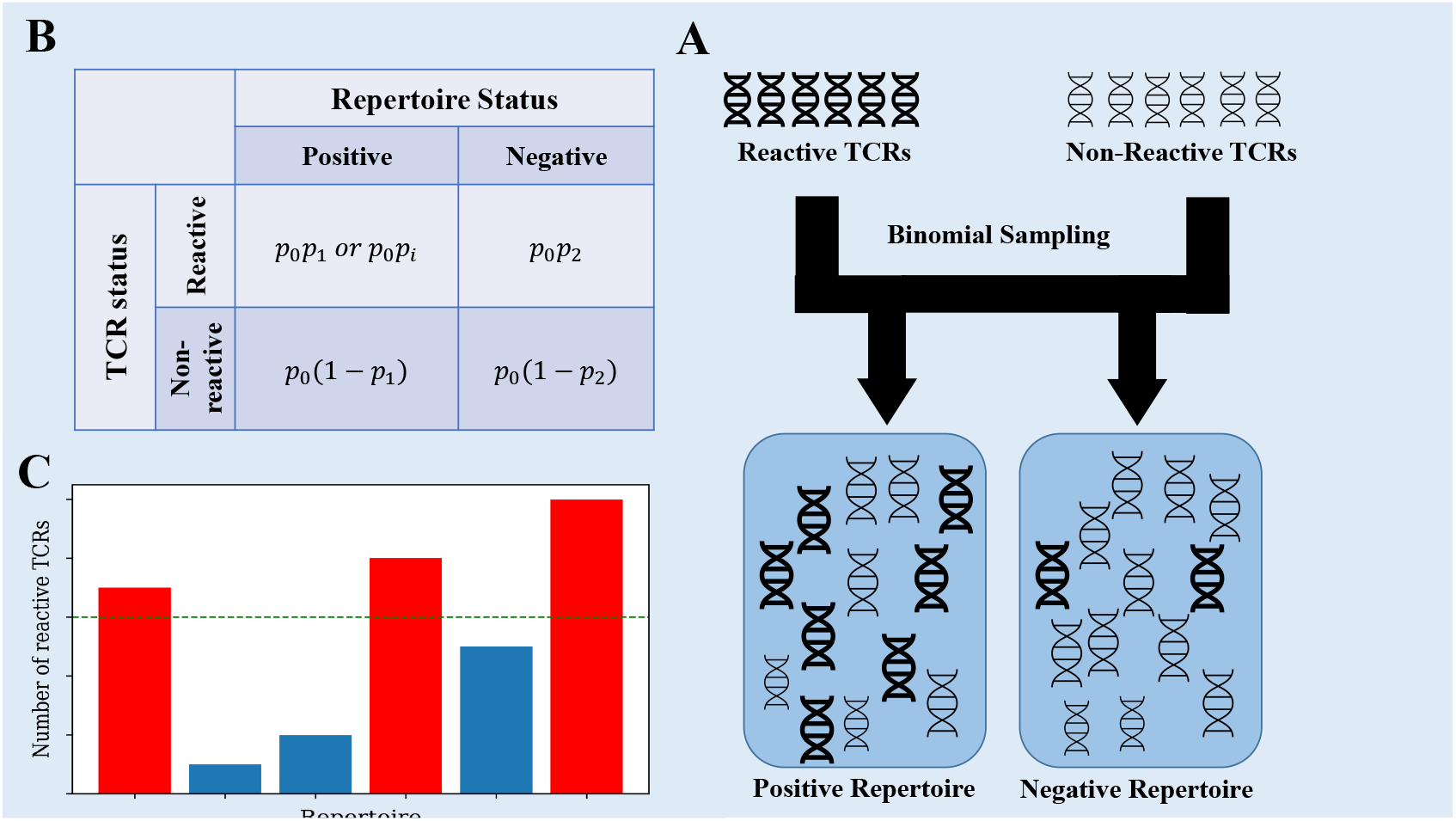
A) Data generation process of the toy model. Each generated repertoire is created using binomial sampling from a collection of positive and negative TCRs. B) The data generation process uses 3 probabilities: *p*_0_ - the probability that a TCR would be selected in any patient, *p*_1_, *p*_2_ - the same for TCRs associated with CMV in CMV+ and CMV-samples. We also tested a model where we replaced *p*_1_ with *p_i_* ~ *N*(*p*_1_, *σ*^2^) for each positive TCR *t_i_*. C) When classifying the generated repertoires, the reactive TCRs are extracted from each repertoire using the *χ*^2^ score on the training set, and then counted in the test set. Repertoires with a large enough number of reactive TCRs are classified as positive.

In this model, all TCRs are independent (the presence or absence of different TCRs are not correlated). In such a model,

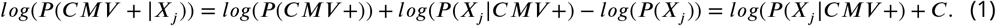

Since the TCRs are independent,

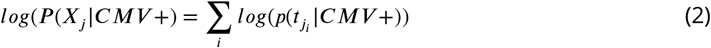

*p*(*t_j_i__*|*CMV*+) are sampled from a binomial distribution. For reactive TCRs *E*(*p*(*x_j_i__*|*CMV*+)) = *p*_0_*p*_1_, whereas *E*(*p*(*x_j_i__*|*CMV*–)) = *p*_0_*p*_2_. Since *p*_2_ ≪ *p*_1_, the negative component can be ignored. Since the *χ*^2^ index awards a high score to TCRs that appear in more positive repertoires than negative TCRs, we can expect that by picking a conservative threshold, most of the TCRs that have an high enough *χ*^2^ are truly reactive (as can be observed from the absence of TCRs with parallele negative scores). However, since general non-reactive TCRs appear in large amounts in both positive and negative repertoires, some might still pass the threshold and be falsely classified as reactive TCRs. When the value of *p*_0_ * *p*_1_ is large enough so that there are much more true reactive TCRs found than false reactive TCRs, we expect that classification to be correct.

We calculated the number of false and true reactive TCRs that are extracted by the *χ*^2^ scoring for different *p*_0_ * *p*_1_ values, using the binomial distribution above (Figure 3A). In the specific sample sizes (see Methods for details of simulations) used here, one can clearly see that by a value of *p*_0_ * *p*_1_ > 0.06 there are considerably more true reactive than false reactive TCRs detected. Below this value, classification would be impossible, while above this value, it should be straightforward. To test that, we applied a straightforward algorithm, where we counted the number of significant TCRs as defined by the training set in each test sample and used the count as a classification score. One can see that the transition between the points that there are more false reactive TCRs than true reactive TCRs to there being orders of magnitude more true reactive TCRs than false reactive TCRs is sharp, and the AUC transition is expected to be similar. As such, either classification is trivial and then counting is enough, or it is impossible and then all other algorithms will also fail. The same holds for all parameter regimes of *p*_2_ and *p*_1_.

**Figure 3.**
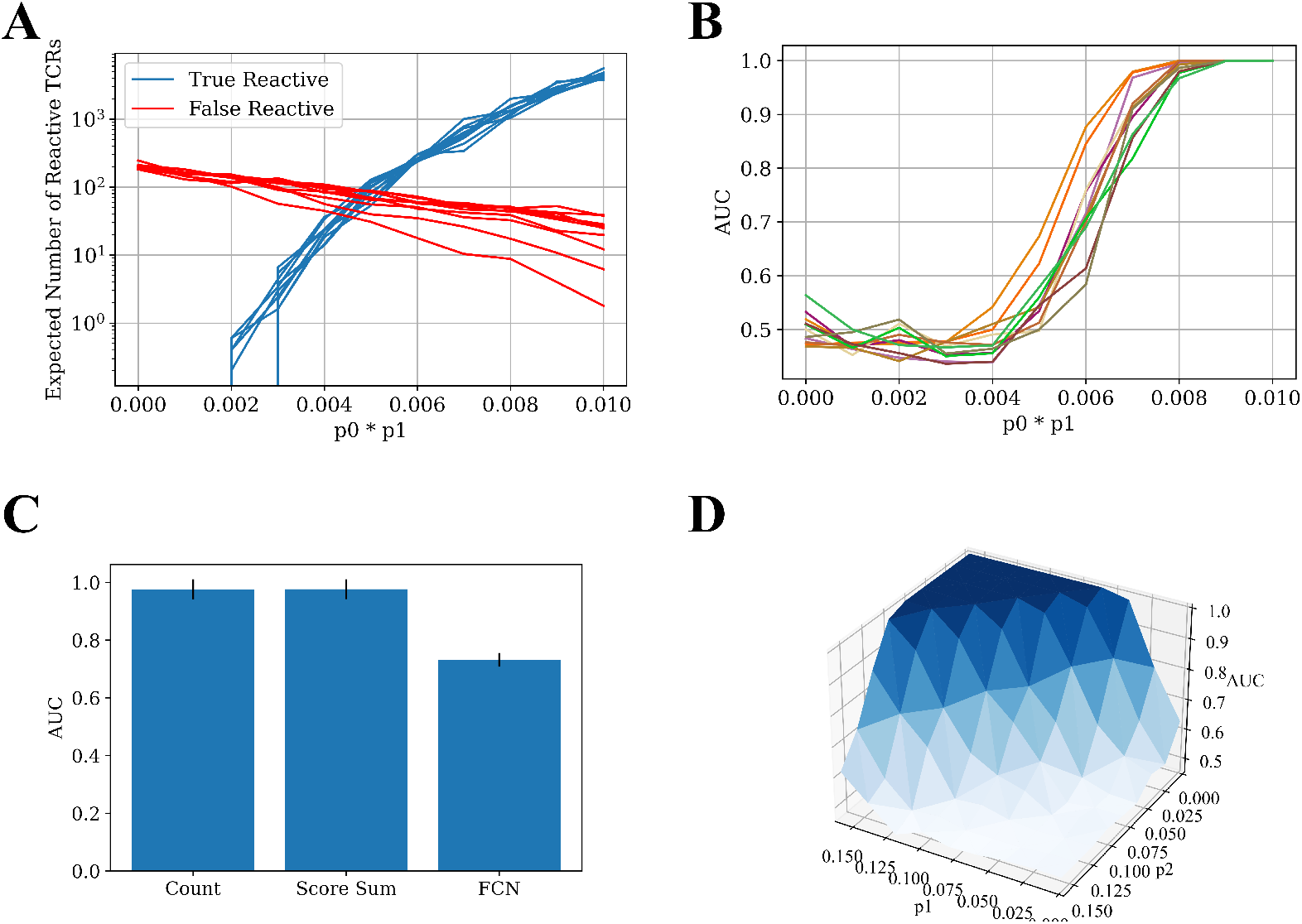
(A) The number of true reactive and false reactive TCRs extracted by the *χ*^2^ scoring. The number is the average of 5 calculations on the training set over a 5 CV splits. Each line represents a constant *p*_0_ ~ *U*[0.01, 0.1] value with different *p*_1_ values. The x-axis is the product of *p*_0_ and *p*_1_. The other parameters are constant: *N* = 100, 000, *p*_2_ = 0.001. (B) The AUC score for data generated with different *p*_0_, *p*_1_ probabilities (5-CV fold). The classification was obtained using the counting method. The colors represents different *p*_0_ ~ *U*[0.01, 0.1] values with different *p*_1_ values. The x-axis is the product of *p*_0_ and *p*_1_. The other generation parameters are as above. (C) Bar plot of the AUC results for different models on the 5 CV above. In all the models, meaningful TCRs are extracted by calculating the *χ*^2^ score for each TCR in the test set, and then taking only TCRs above a certain threshold (in this case, 3.84). The counting model counts the relevant TCRs in each test set sample and classifies it by the number of relevant TCRs in each repertoire. The score sum model sums the *χ*^2^ score for the relevant TCRs in the test repertoires, and classifies them according to the sum. The FCN model trains a 2-layer FCN over the training repertoires and then makes a prediction on the test repertoires using the TCR one-hot vectors as an input. The parameters used in the generation of the repertoires are *N* = 100,000, *p*_0_ = 0.1, *p*_1_ = 0.08, *p*_2_ = 0.002. (D) A surface plot that presents the AUC of the counting model for different *p*_1_ and *p*_2_ combinations. Here, *p*_1_ is not constant for each TCR. Instead, *p_i_* is sampled from for each TCR *t_i_* (see Figure 2) from a normal distribution. The other generation parameters are constant: *p*_0_ = 0.01, *σ* = 0.03, *N* = 100,000.

The test generated data (500 positive repertoires, 500 negative repertoires) and was split into a test and a training set. Reactive TCRs were extracted from the training set, and counted in each sample in the test set. Then, an AUC score was calculated using the number of positive clones present in each repertoire in the test set. We ran the counting model on the generated data with different parameters. As expected from the argument above, when trying to classify the generated data with a low value of *p*_0_ * *p*_1_, the classification is impossible. With a high enough value of *p*_0_ * *p*_1_, the classification is almost trivial, and a simple counting model can achieve a perfect AUC (Figure 3B). More importantly, the range between the two extremities is very narrow, either you can or cannot classify the repertoires using counting. Since there is no a priori reason to assume for any disease and sampling level in any given experiment that they are exactly in this narrow range, one can argue that in general for any disease, either classification is impossible, or a simple counting argument can obtain a high accuracy.

Given this simple argument, one would expect other methods to simply overfit in the simulation above. To test for that, we compared the counting with more complex methods (see Methods). Indeed, counting the relevant TCRs is the best repertoire classification method. The introduction of machine learning methods often only reduce the classification accuracy, following over-fitting on the training set (Figure 3C).

To ensure that the results are not an artifact of the highly simplified model, where all the positive TCR have the same probability, we further enlarged the model to contain a different a priori probability for each positive TCR to appear (see Methods). Figure 3D shows that the conclusion of the sharp transition is true even with looser conditions. Even when *p*_2_ is changing, and when the reactive TCRs are sampled in a non constant distribution, there is still a clear and sharp “tipping point” between impossible and easy classification, suggesting that this argument may apply to real sampled data.

### Application to real data

To show that the counting argument works in general even when not all TCRs are independent, we analyzed the immune repertoire ECD ((15)).

To test for the CMV classification, we split the data into a training:validation:test split ratio of 8:1:1, and used 9 cross validations on the training and validation (the test set was either not changed or ever used in the training). We then applied the counting method:

1. Calculate the *χ*^2^ score for each TCR in the training set.
2. Extract the top-*k* TCRs with the highest *χ*^2^ score. In this case *k* = 100. One could alternatively use a *p* value cutoff with similar values, but we have here tried to minimize the hyperparameter optimization to show how generic the counting algorithm is.
3. Count the number of reactive TCRs in each test sample.
4. Calculate AUC on the test set using the counts above.

Again, the counting model outperformed all published models, including the (15) model on the same test set for different training set sizes (Figure 6). The advantage of the counting algorithm is further obvious in small training sample sizes. In contrast with Emerson (15) and deepRC (52), the counting method can obtain a signal even for 100 training samples.

**Figure 4.**
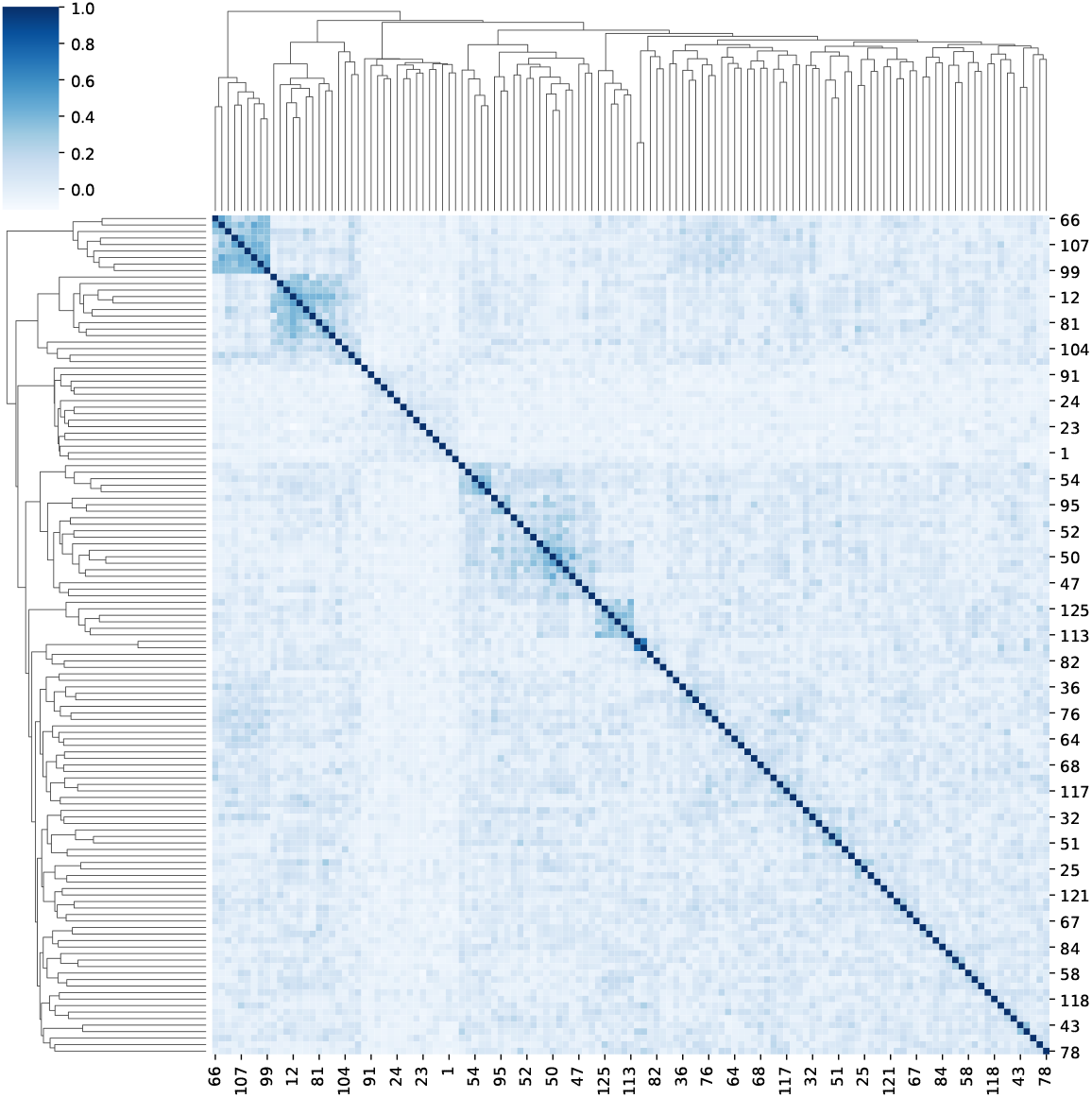
A clustermap of the Spearman correlation between 125 reactive TCRs. For each reactive TCR, we extracted from the ECD (15), we assigned a one-hot vector that represents the appearance on the TCR in different repertoires in the data. Then, for each TCR pairing, we calculated the Spearman correlation between their one-hot vectors.

**Figure 5.**
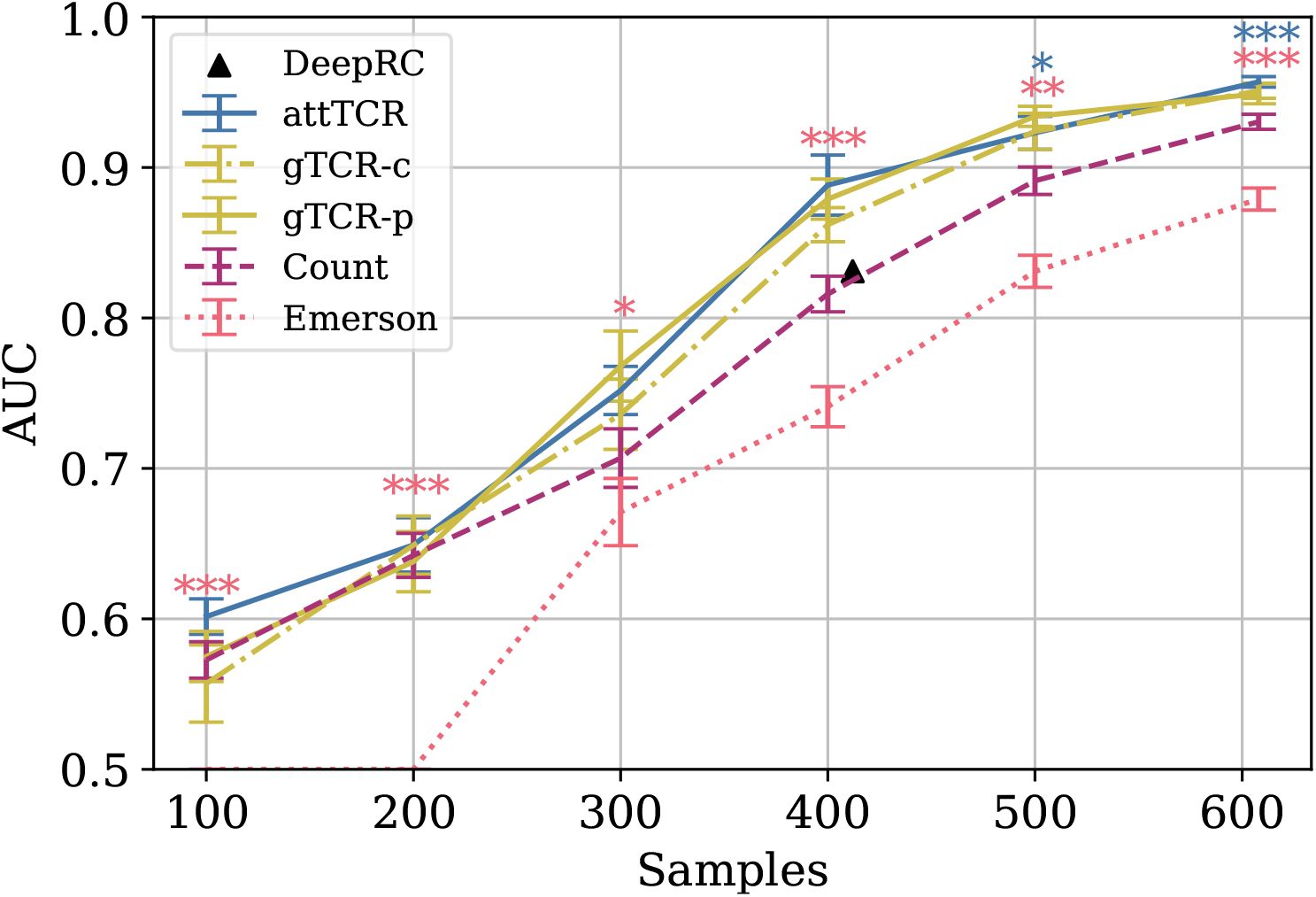
The AUC results of different models on different train sample sizes on the ECD (15). The results are over a 9 CV split of the training and the dev sets. The test set is the same for every model. Stars are used to mark statistical significance of the results using a t-test (* *p* < 0.05, ** *p* < 0.01, *** *p* < 0.001). Pink stars represent the t-test between the Emerson model and the counting model, and blue stars represent the t-test between attTCR and the counting model. The results were also compared to the results reported for deepRC ((52)) (with different experimental setup). For further result comparison to DeepRC and MotifBoost on the ECD, which have even lower AUC, see (28)

**Figure 6.**
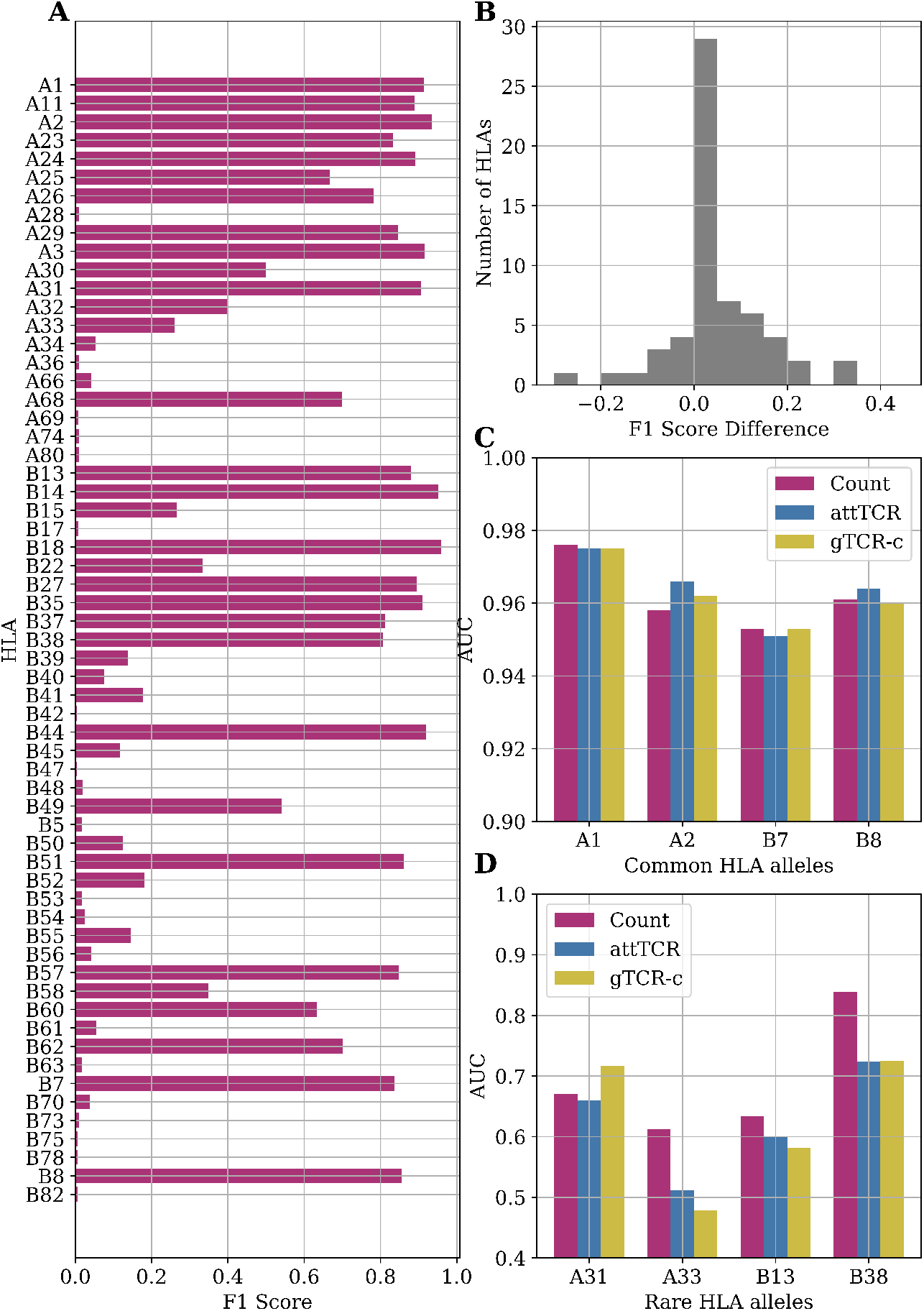
A) F1 score results for the counting model on HLA classification. We performed a leave one out split over the entire dataset. B) A histogram of the F1 score differences between the classification results of the counting model and the Emerson model. The difference between the counting model and the Emerson model is statistically significant (*p* = 0.017 with Mann-Whitney U-test) (15) on the same HLA alleles. C) Comparison of AUC results of the counting model, attTCR and gTCR-c on the repertoire HLA classification task. A 5-fold CV was used, and the AUC was calculated using prediction pooling instead of averaging (6). The HLA alleles presented are the most frequent HLA alleles in the dataset. D) Comparison of AUC results of the counting model, attTCR and gTCR-c on the repertoire HLA classification task. The HLA alleles presented are the least frequent in the dataset.

### TCRs Correlations

In contrast with the simplistic model, TCR usage in real samples can be correlated. The counting method, as adequate as it is, neglects the information that may be available in this correlation. As such it does not reach a perfect AUC in the ECD. To check the co-expression of reactive TCRs, we computed the Spearman correlation between the appearance vector of each TCR in each sample (1 if the TCR is in the sample and 0 otherwise (Figure 4), and clustered the samples based on their correlation). The clusters of related TCRs are very clear. To test the significance, we used a t-test between the correlation matrix and a correlation matrix between random shuffled vectors (*p* < 1.*e* – 100).

### Autoencoder Projections

To address the similarity between TCRs, one can use either a sequence similarity (how similar are the TCR CDR3 and V sequence), or a functional similarity (how often they co-appear in the same sample). For the sequence similarity, we projected each sequence using an improvement of the ELATE (Encoder based LocAl Tcr dEnsity) TCR autoencoder (13). ELATE was enlarged to become a cyclic variational autoencoder, and the TCR representation method was improved (see Methods).

To confirm that the autoencoder projection is associated with the class of the TCRs, we sampled 100 TCRs out of the 200 TCRs with the highest *χ*^2^ score, and 100 random TCRs, and computed the average nearest neighbor euclidean distance between the projections within each group (with 30 cross validations). The distance between reactive TCRs is significantly lower than random TCRs (12.95 vs 13.886, T test *p* < 1.*e* – 10), suggesting that reactive TCRs are evenly distributed among all TCRs.

### attTCR

In order to combine the projections into a classifier, we propose an attention model. However, classical attention models sum the positive attention scores to 1. As such, these models would fail to count the number of reactive TCRs in a sample. Instead, they would focus on the relative importance of reactive TCRs. We thus propose a novel attention model that does not apply a softmax to the score assigned to each reactive TCR (see Methods for details), but sigmoid. As such it allows to estimate the relative importance of reactive TCRs and on the other hand to count them. The sum is then normalized to be in the active range for the loss function. We then tested the combination of the projection and the attention on the ECD, and the results are significantly better than the counting algorithm (Figure 5, for every training set size, and *p* values of differences therein), and obviously much better than all existing models.

### gTCR

attTCR has an impressive precision. However, it its complex and its training is costly (in GPU time). An alternative method to incorporate the relation between TCR would be purely based on their co-occurrence in samples. To address that, we propose a novel GNN formalism that we denote Graph TCR (gTCR). We define a graph connecting TCRs based on the correlation between their cooccurrence patterns (two TCRs are connected if the Spearman correlation coefficient between their co-occurrence vector is above 0.2). Then the occurrence vector of each TCR in a given sample is the input of this GNN In parallel, the log frequency of each TCR is included as the input to an FCN and the last layers of the two are the input of a final FCN layer that combines the interaction map with the co-occurence and the log frequency. The results of gTCR are close to the results of attTCR, with no significant difference (Figure 6). Note that similar results can be obtained by producing a graph using the similarity of the TCRs projection (denoted in the figure gTCR-p in contrast with gTCR-c).

The difference between the two gTCR models is simply the interaction matrix, which can be either based on the sequence or the appearance similarity. The resulting interaction matrices are very different (Jaccard index = 0.002 ± 0.003 in 10 training/test division). Thus, information seems to be available through both distance definitions.

### HLA allele repertoire classification

The ECD ((15)) provides the low resolution A and B HLA-alleles of most samples. We further tested the algorithms above HLA prediction accuracy. From a MIL point of view, this is equivalent to CMV classification. Indeed, the counting model handles this classification task very well, especially with very frequent HLA alleles (Figure 7A). The difference between the counting model and the Emerson model is statistically significant (*p* = 0.017 with Mann-Whitney U-test). We use the counting model with *k* = 100. We use a number cutoff instead of a threshold cutoff to ensure that we find reactive TCRs for rare HLA alleles. Those TCRs receive a relatively low *Chi*^2^ score to the reactive TCRs since very few samples have them.

**Figure 7.**
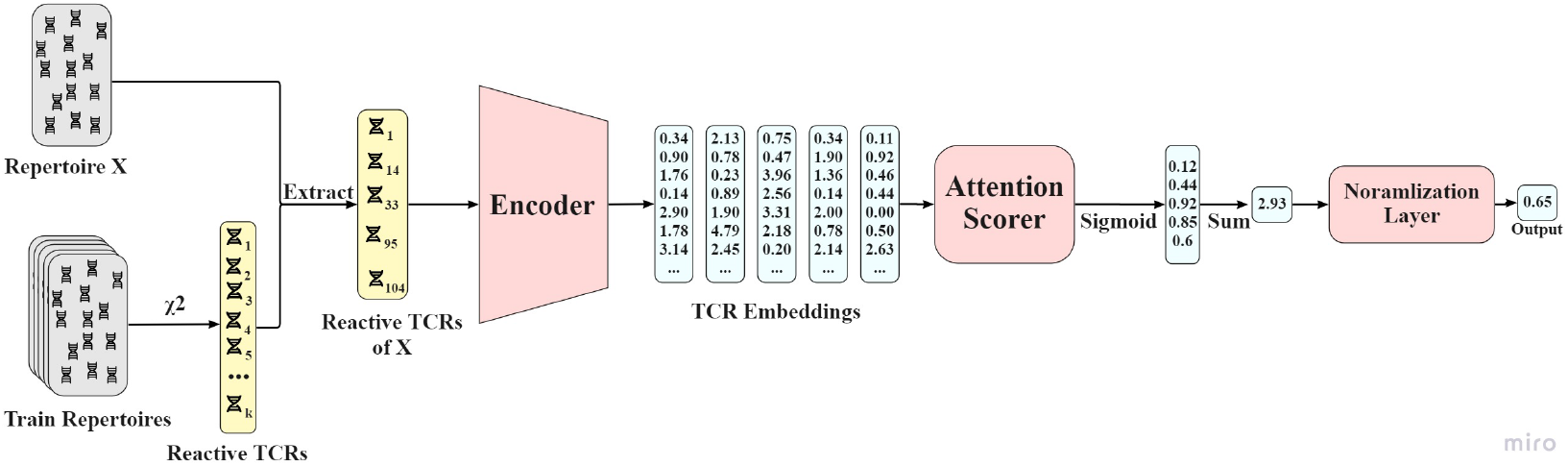
AttTCR’s architecture. First, the reactive TCRs are sampled from all the train repertoires using the *χ*^2^ method. Then, for each repertoire *X*, the reactive TCRs contained in *X* are extracted. Each reactive TCR is projected by the encoder. The projections are then scored by the attention scorer. The scores are summed and normalized. The output of the model in a number between 0 and 1 that indicates the confidence of the model on whether the repertoire is positive.

The counting model has a higher accuracy than the Emerson model on most HLA alleles (Figure 7B). Machine learning models, specifically attTCR and gTCR-c, have similar results to the counting model for common HLA alleles (Figure 7C), but over-fit for rare HLA alleles (Figure 7D). For full results over all the HLA alleles, see the Appendix.

Multiple other comparisons were proposed, such as taking the MIRA(nol) Covid-19 samples as positive repertoires and the ECD as negative repertoires (since there was no COVID-19 at the sampling time), and the counting method obtains an AUC of 1 on this comparison. However, this may be a batch effect, since the two samples may have have differences in the sampling and analysis protocol.

## Methods

### Simulated samples

In order to analyze the performance of the classification methods, we propose a simple simulation that captures the essence of the classification problem. Assume a general very large set of TCRs, where each patient has a random subset of these TCRs. Within the large set of TCRs, there is a small subset associated with the disease, and patients that had the disease have a higher than random chance of having these TCRs (see Figure 2 for description of model). The data generation process uses 3 probabilities: *p*_0_ - the probability that a TCR would be selected in any patient, *p*_1_, *p*_2_ - the probability that a chosen TCR is associated with positive and negative samples. We also tested a model where we replaced *p*_1_ with *p_i_* ~ *N*(*p*_1_, *σ*^2^) for each reactive TCR *t_i_*.

In the different trials performed in the current analysis, we generated 1,000 different repertoires (500 positive, 500 negative) using differing generation probabilities (*p*_0_, *p*_1_, *p*_2_). All the experiments were performed using a 4:1 training:test split, using 5 different data splits.

### *χ*^2^ Score

To extract reactive TCRs from a repertoire, we use a simple scoring method. For each TCR *t*, the *χ^2^* formula uses the following values:

- *N pos_i_* - The number of positive repertoires that contain *t_i_*.
- *N_i_* - The total number or repertoires that contain *t_i_*.
- *N pos* - The total number of positive repertoires in the data.
- *N* - The total number of repertoires in the data.

The *χ^2^* score for TCR *t_i_* is calculated using Equation 3.

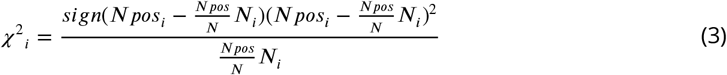

The difference with the regular *χ*^2^ is simply the sign of the deviation.

### Counting Model

The counting model is a simple model that effectively manages to distinguish between positive and negative repertoires on the test set. The counting model has the following steps:

1. Calculate the *χ*^2^ score for each TCR in the training set.
2. Extract all the TCRs with a *χ*^2^ score over a certain threshold. The threshold can be either a *p*-value, or a fixed number of TCRs *k*.
3. Count the number of significant reactive TCRs in each file of the test set.
4. Calculate AUC on the test set using the counts.

### TCR Autoencoder

A TCR autoencoder is a model that preserves the information about input *t_i_* V gene and CDR3 sequence, while reducing the dimension to a low dimension representation *z_i_*. The training of the TCR autoencoder includes several steps of data processing (13). The first step is representing each of the amino acid per position as well as the V genes by an embedding vector. There are twenty possible amino acids and an additional end signal is required. Each instance is then processed by an autoencoder network and encoded to size 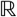^30^ (we have previously checked that adding dimensions beyond 30 had a very limited contribution to the accuracy).

The autoencoder network contains three layers of 800, 1100, and 30 neurons as the encoder and a mirrored network as the decoder. The network is trained with a dropout of 0.2 and a ReLU. An MSE loss function is implemented to compare each input sequence to the resulting decoded sequence (13). The current version differs from the ELATE encoder ((13)), since it includes a variational term. Instead of encoding an input as a single point, we encode it as a distribution over the latent space. The model is then trained as follows: First, the input is encoded as a distribution over the latent space;second, a point from the latent space is sampled from that distribution, Then the sampled point is decoded and the reconstruction error can be computed;finally, the reconstruction error is back-propagated through the network. The VAE loss function is the same as ELATE with a Kulback-Leibler divergence between the returned distribution and a standard Gaussian.

The problem with the standard VAE is that the KL term tends to vanish. A recent work ((17)) studied scheduling schemes for *β*, and showed that KL vanishing is caused by the lack of good latent codes in training the decoder at the beginning of optimization. To remedy this, we used a cyclical annealing schedule, which repeats the process of increasing *β* multiple times. This new procedure allows the progressive learning of more meaningful latent codes, by leveraging the informative representations of previous cycles as warm restart.

### attTCR

The attention model receives as an input the reactive TCRs of each repertoire, and as an output a score between 0 and 1 that predicts whether the repertoire is positive or negative. The model is composed of an encoder network, an attention scorer, and a normalization layer. The encoder was explained above.

#### Attention Network

Each TCR *t_i_* is assigned an attention score *a_i_*, such that *a_i_* ∈ [0, 1]. TCRs that are more important to the classification should receive higher attention scores. The attention network takes as an input the embedding of each TCR by the encoder network and is composed of 2 hidden layers of size *q*. The output of the attention network for each TCR sequence is a single attention score. Therefore, for the entire repertoire, the network outputs a vector *ν* of dimension *N* (the number of reactive TCRs). A sigmoid function is used to produce an attention score between 0 and 1 for each reactive TCR in the repertoire. We have here used the traditional Transformer ((49)) notation. We use the following matrices and vectors to describe the attention process:

- 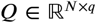 - The queries matrix. In our model, the matrix is created after the 2 hidden layers of the attention network.
- 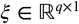 - The keys vector. The weights of the output layer of the attention network.

The attention score calculation applied by equation 4, where *σ* is the sigmoid function:

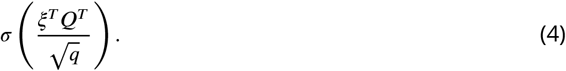

Note that unlike traditional attention models, we do not use the softmax function on the resulting attention vector, nor do we multiply each attention by the TCR representation. We are not interested in performing a weighted average. Instead, we want to score each TCR and still keep the information about the number of reactive TCRs in the repertoire, i.e., *N*. Thus the score of an entire repertoire is simply the sum of the attention values for all the reactive TCRs in this sample.

#### Normalization Layer

The input of the normalization layer in a vector 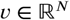 with the scores of each TCR in the repertoire. The Normalization layer’s goal is to convert the sum over the vector *v* to a number between 0 and 1, so we can train the model using BCE loss. Just putting **∑** *v* into a sigmoid function is not going to work, since the sum of *N* scores *a_i_* ∈ [0, 1] is very likely to be too large for the sigmoid function. As a result, all the repertoires would output a number very close to 1, which might hurt the training process. Therefore, we use 2 learned parameters: *γ*_1_, *γ*_2_, to normalize the sum before the sigmoid function. In conclusion, the normalization layers performs Equation 5.

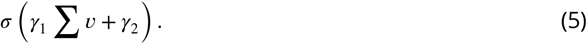

### gTCR

#### Graphs

##### TCR similarity graph

We define here two ways of modeling the TCR-graph. Both ways consists of two stages, a definition of similarity matrix between the reactive TCRs followed by a zeroing stage where rows and columns from the similarity matrix are filled with zero value if the reactive TCR is absent from the sample’s repertoire.

One way of modeling such a similarity matrix between reactive TCRs is obtained using the Spear-man correlation matrix between the training samples presence vectors. These sample’s presence vectors contain 0 or 1 according to the presence of each reactive TCR in the sample’s repertoire. Another way of modeling a similarity matrix between reactive TCRs is obtained using the inverse of the euclidean distance between the projection of the reactive TCRs obtained from the autoen-coder.

##### gTCR

The gTCR (graph TCR) model combines the information from the frequencies vector as well as the graph represented by the normalized adjacency matrix as can be seen in Equation 6. An embedding vector of the log frequencies is obtained from a 2-layer FCN, each followed by a tanh activation function and dropout layer (Equation 8). In parallel, one layer of a GCN model is applied (Equation 9) to the reactive TCR presence vector. Then the output of the two networks are concatenated and serve as the input of a 2-layer FCN to predict a binary condition (Equation 10).

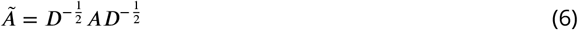

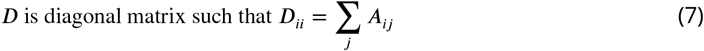

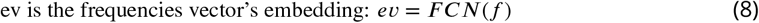

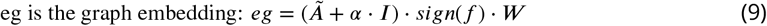

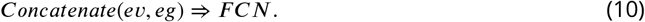

*α* is a learned scalar regulating the importance given to the vertex’s feature compared to its neighbors features. *α* is initialized with 1 plus Gaussian *N*(0, 0.1).

### Comparison to Other Methods

The counting method was compared to 2 other classification methods:

- **Score Sum** - This method is almost entirely similar to the counting model. The only difference is that instead of classifying the repertoires based on the number of reactive TCRs found in the repertoires, we classify them by the sum of the *χ*^2^ scores of the reactive TCRs in each repertoire.
- **FCN** - After extracting the reactive TCRs from the data, each repertoire is embedded to a vector of the dimension of the number of reactive TCRs. Each dimension in the vector represents a different reactive TCR, and its value is set to 1 if the repertoire contains the TCR and 0 otherwise. Then, a 2-layer FCN is fitted on the vector training set, and tested on the test set.

### Data

The Emerson dataset contains 786 immune repertoires (15). Each repertoire contains between 4,371 to 973,081 (avg. 299,319) TCR sequences with a length of 1 to 27 (avg. 14.5) amino acids. The V and J genes and the frequency are saved for each TCR. 340 repertoires are labeled CMV+, 421 are labeled CMV-, and 25 are of unknown status. We only use the repertoire with a known CMV status, 761 repertoires in total. In addition 626 of the repertoires have HLA allele information available.

### Preprocessing

The Emerson dataset (15) is composed of 786 repertoires in total. However, since the task at hand is a supervised classification task, the 25 repertoires without a CMV classification are not beneficial to the learning process, so they are removed from the dataset. Then, all the TCRs that have missing CDR3 amino acid information are discarded. In the following step, the TCRs are filtered based on prevalence in different repertoires. Only TCR sequences that appear in 7 different repertoires or more remain in the repertoires after the filtration.

### Experimental Setup

When predicting CMV status of the repertoires, the models are tested with a test size that contains 10% (77 samples) of the data. For all the models tested, the test set is the same. attTCR is trained using a 9-fold CV between the training set and the validation set, while gTCR is trained over 20 different splits. In the counting model, the validation set is not used. All the models are evaluated using an AUC score on the test set (31).

The HLA allele repertoire classification in Figure 7A was evaluated using an F1 score. In Figures 7C and 7D, the measure was changed to AUC over a 5-fold CV with a train:validation:test split of 3:1:1. The AUC was calculated using the pooling method, i.e., calculated once over all the predictions (6).

## Discussion

We have here proposed three novel methods with different levels of complexity, and shown that even the simplest of these models outperform the current State of The Art (SOTA) for repertoire classification. The simplest model is simply counting reactive TCRs, followed by a novel attention model that combines classical attention models with counting, and finally a combination of graph based machine learning with MIL. All the models presented in the paper rely on the assumption that TCRs are only positively selected, and there are no TCRs negatively associated with a condition. Note that (15) used a Fisher exact test to score TCRs based on their association with positive and negative repertoires. It also classifies each significant TCR as either a positive or a negative selected TCR. However, the assumption that there are any negatively selected TCRs does not make much immunological sense. TCR expansion occurs when a certain TCR binds to an antigen-peptide. There is no equivalent process for TCRs that do not bind to antigen-peptides. Thus, in theory, the abundance of a TCR in a repertoire can only indicate that the TCR was positively selected.

Some TCRs are highly abundant in different individuals (25), and have initial production probability (14; 3). Therefore, positively selected TCRs exist in various frequencies in positive immune repertoires, some especially common ones might appear randomly in negative instances as well. Thus, the relative abundance of a TCR in many repertoires does not automatically make it more indicative than a TCR that appears in a few repertoires. Once a TCR is proven to be positively selected, its frequency does not matter much when it comes to repertoire classification. Hence, the counting model is a good way to classify the repertoires given the reactive TCRs.

We have shown in the data that TCRs are indeed only positively selected, and that it improves on existing models in both theory and real data. There are distinctions to be made between the real repertoire data, and the generated one. The most obvious is that real TCR presence in a repertoire does not follow a binomial distribution. Real TCRs have a scale free distribution. Some TCRs are public TCRs and are very common (25), and others are very rare. The pool size of positive and negative TCRs to draw from is also vastly different in size. Statistically, there are many more possible negative TCRs than TCRs that bind to a epitope-peptide of a specific disease. In addition, TCRs are sampled in varying sizes, whereas the repertoires in the generative model are all around *P_0_N*. Despite these differences, we believe that the conclusions of the toy model are still true on real repertoire data. However, these differences do not affect the validity of counting and its extension.

The current approach is purely based on the observed TCR presence and absence and on their sequence. It completely ignores the antigen or MHC properties. However, multiple algorithms were proposed for both TCR-peptide (21; 10; 44; 27; 37; 42; 14; 16; 19; 45; 35; 47; 5; 26; 11) and TCR-MHC binding (22; 54; 32; 2; 34; 38; 40; 29; 50; 20; 51; 30). While the accuracy of such algorithms keeps improving, it is still too, it may be too early to use such algorithms for repertoire classification.

The ML models presented in the paper, especially attTCR, can also be used in a large variety of problems. attTCR presents a new approach of attention scoring, that can be used in every MIL task that involves counting. Further research has to be done on the quality of the proposed ML models on other non-related tasks. However, we propose that these three levels of modeling - counting, counting attentions models and GNNs on selected shared samples may be a general approach to all MIL problems.

## Appendix 1

In the paper, we have shown that repertoires can be classified using bayesian and machine learning tools on the content of the TCR repertoire. However, in the Appendix we want to prove that there does not exist an easier, more superficial method of distinguishing between positive and negative repertoires. In Figures 1 and 2 we show that different general attributes of the repertoires are the same with positive and negative repertoires, and they cannot be differentiated using this attributes.

**Appendix 1 Figure 1.**
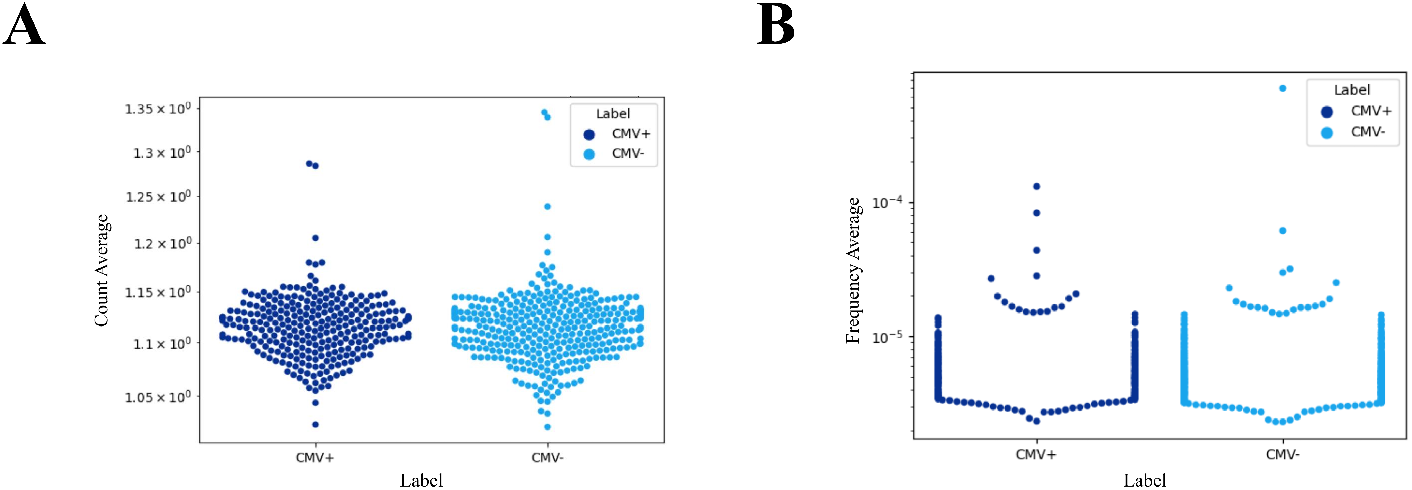
(A) A swarm plot of the different repertoires in the data. Each dot represents a repertoire. The y-axis represents the average count of a TCR in a repertoire, where count of a TCR is defined as the number of clones the TCR has in the repertoire. It is clear that there is not a big difference in the count distribution between positive and negative repertoires. (B) A swarm plot of the different repertoires in the data. Each dot represents a repertoire. The y-axis represents the average frequency of a TCR in a repertoire. It is clear that there is not a big difference in the frequency distribution between positive and negative repertoires.

**Appendix 1 Figure 2.**
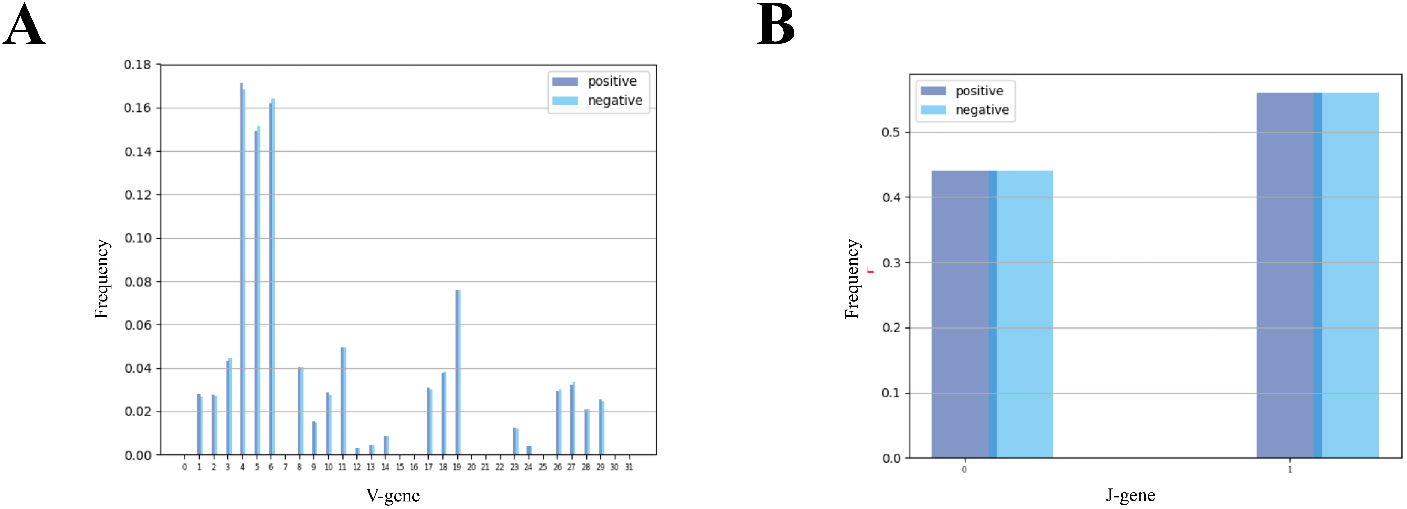
(A) A histogram of the different *Vβ*-genes in the data. Each column represents the average frequency of a *Vβ*-gene in positive and negative repertoires. It is clear that the v-gene distribution between negative and positive repertoires is very similar. (B) A histogram of the different *Jβ*-genes in the data. Each column represents the average frequency of a *Jβ*-gene in positive and negative repertoires. It is clear that the J-gene distribution between negative and positive repertoires is very similar.

